# Dynamic culture of cerebral organoids using a pillar/perfusion plate for the assessment of developmental neurotoxicity

**DOI:** 10.1101/2024.03.11.584506

**Authors:** Prabha Acharya, Sunil Shrestha, Pranav Joshi, Na Young Choi, Vinod Kumar Reddy Lekkala, Soo-Yeon Kang, Gabriel Ni, Moo-Yeal Lee

**Affiliations:** Department of Biomedical Engineering, University of North Texas, Denton, Texas; Bioprinting Laboratories Inc., Dallas, Texas

**Author notes:** Corresponding author: Moo-Yeal Lee.

**Keywords:** Dynamic organoid culture, Pillar plate, Perfusion plate, Cerebral organoids, Developmental neurotoxicity

## Abstract

Despite the potential toxicity of commercial chemicals to the development of the nervous system (known as developmental neurotoxicity or DNT), conventional *in vitro* cell models have primarily been employed for the assessment of acute neuronal toxicity. On the other hand, animal models used for the assessment of DNT are not physiologically relevant due to the heterogenic difference between humans and animals. In addition, animal models are low-throughput, time-consuming, expensive, and ethically questionable. Recently, human brain organoids have emerged as a promising alternative to assess the detrimental effects of chemicals on the developing brain. However, conventional organoid culture systems have several technical limitations including low throughput, lack of reproducibility, insufficient maturity of organoids, and the formation of the necrotic core due to limited diffusion of nutrients and oxygen. To address these issues and establish predictive DNT models, cerebral organoids were differentiated in a dynamic condition in a unique pillar/perfusion plate, which were exposed to test compounds to evaluate DNT potential. The pillar/perfusion plate facilitated uniform, dynamic culture of cerebral organoids with improved proliferation and maturity by rapid, bidirectional flow generated on a digital rocker. Day 9 cerebral organoids in the pillar/perfusion plate were exposed to ascorbic acid (DNT negative) and methylmercury (DNT positive) in a dynamic condition for 1 and 3 weeks, and changes in organoid morphology and neural gene expression were measured to determine DNT potential. As expected, ascorbic acid didn’t induce any changes in organoid morphology and neural gene expression. However, exposure of day 9 cerebral organoids to methylmercury resulted in significant changes in organoid morphology and neural gene expression. Interestingly, methylmercury did not induce adverse changes in cerebral organoids in a static condition, thus highlighting the importance of dynamic organoid culture in DNT assessment.

## Introduction

Exposure to chemicals during pregnancy has the potential to adversely influence the normal development of the human brain, a phenomenon known as developmental neurotoxicity (DNT) of compounds ^1^. The developing brain is more susceptible to chemical exposure compared to the adult brain because the protective blood-brain barrier is not fully functional in embryo, fetus, and newborn ^2–7^. The brain development process including differentiation, proliferation, migration, and cell-cell communication must take place in a controlled time frame, and any disturbance in this process by a trace amount of chemical exposure could lead to permanent damage to the developing brain ^8^. Despite the potential vulnerability of the developing brain to compounds, only a few hundred compounds among tens of thousands of chemicals have been tested for DNT. Conventional DNT assessments rely entirely on animal testing, a complex and challenging process with limited applicability to subsequent human trials ^1,9,10^. As a result, there is an urgent need for reliable, rapid, and economical *in vitro* screening systems that can mimic human-specific developmental pathways to evaluate the DNT potential of chemicals. Conventionally, *in vitro* studies are mostly based on short-term cytotoxicity assays and do not focus on long-term changes in cell differentiation, morphology, and gene expression relevant to brain development ^8,11,12^. In addition, the majority of neurotoxic compound screening have been conducted on two-dimensional (2D) cell monolayers, which lack the physiological relevance to accurately recapitulate brain development processes *in vivo* due to their limited ability to replicate essential cell-cell and cell-matrix interactions ^13,14^.

Recently, neurospheres and brain organoids have been utilized for evaluating the DNT potential of compounds by closely mimicking *in vivo* physiology ^11,12,15,16^. Notably, brain organoids differentiated from induced pluripotent stem cells (iPSCs) have opened new avenues for studying human brain development, function, and disease pathways at cellular and structural levels ^17–22^. Several published iPSC-derived brain organoid differentiation protocols have potential to mimic the normal brain development ^23–25^. While human brain organoids offer a unique and innovative approach for evaluating DNT potential of chemicals in predictive *in vitro* screening, there remain several technical challenges that need to be addressed before adopting them in chemical product development. Existing brain organoid culture systems, such as 6/24-well plates, petri dishes, and spinner flasks, requiring a substantial volume of expensive culture media, are not amenable to high-throughput compound testing and organoid imaging ^26^. Another significant issue is the batch-to-batch variation in brain organoid culture, mainly due to long-term (typically 1 to 6 months) cell differentiation in hydrogels required to enhance organoid maturity. In addition, due to the prolonged cell culture required, the size of brain organoids often exceeds the diffusion limitation of nutrients and oxygen, leading to the formation of the necrotic core ^27–29^. To address the diffusion issue, microphysiological systems (MPS) have been utilized for brain organoid culture, striving to replicate shear stress and blood flow as found *in vivo*, thereby enhancing organoid maturity and complexity ^29–33^. Nonetheless, current microfluidic devices suffer from low throughput and user unfriendliness due to pumps and tubes required for operation.

To fully utilize the potential of brain organoids for predictive DNT assessment, there is an urgent need for miniature, high-throughput, microfluidic systems that provide the environments for dynamic organoid culture and allow *in situ* compound testing ^22^. To address this need, we have developed a unique pillar/perfusion plate platform and demonstrated robust human organoid culture in static and dynamic conditions ^34,35^. In the present work, we reproducibly generated human cerebral organoids in the pillar/perfusion plate by transferring embryonic bodies (EBs) in an ultralow attachment (ULA) well plate to the pillar plate and differentiating them in the perfusion plate for 1 – 2 months. A small volume of iPSC differentiation media was added in the reservoirs of the perfusion plate on a digital rocker, which generated bidirectional flow in the perfusion wells where the pillars with EBs were inserted for long-term, dynamic cerebral organoid culture, without using any pumps and tubes. Dynamically cultured cerebral organoids showed enhanced viability and maturity with no apparent necrotic core, as compared to their counterpart, statically cultured cerebral organoids. An early stage of cerebral organoids in the pillar/perfusion plate were differentiated with model compounds for 1 and 3 weeks to assess DNT potential. We envision that the pillar/perfusion plate holds promise for organoid-based, high-throughput, predictive compound screening due to its miniaturization, user friendliness, and ability to perform *in situ* organoid imaging.

## Materials and Methods

### Design of the pillar/perfusion plate for dynamic organoid culture

A 36-pillar plate with sidewalls and slits (“36PillarPlate”) and a 36-perfusion well plate with reservoirs and microchannels (“36PerfusionPlate”) were designed with SolidWorks software (SolidWorks Research Standard and Simulation Premium 2022, Waltham, MA, USA) for dynamic organoid culture (**Fig. 1A**). The simulation of flow patterns within the perfusion wells has been performed for optimal dynamic cell culture ^34,35^. Both 36PillarPlate and 36PerfusionPlate were manufactured by injection molding with polystyrene and were functionalized with poly(maleic anhydride-*alt*-1-octadecene) and alginate (Bioprinting Laboratories Inc., Dallas, TX, USA) ^34,35^. The 36PillarPlate with a 6 x 6 array of pillars (4.5 mm pillar-to-pillar distance, 11.6 mm pillar height, and 2.5 mm outer and 1.5 mm inner diameter of pillars) was used for spheroid/organoid culture within the hydrogel scaffold. The complementary 36PerfusionPlate has a 6 x 6 array of perfusion wells (3.4, 3.4, and 11.9 mm well width, length and depth, and 4.5 mm well-to-well distance), a 2 x 6 array of reservoirs (3.6, 20.1, and 15.4 mm reservoir width, length and depth, and 4.5 mm reservoir-to-reservoir distance), and microchannels connecting perfusion wells and reservoirs (**Fig. 1B**). Thus, there are six fluidic channels in the 36PerfusionPlate, each channel composed of six perfusion wells connected by microchannels between two reservoirs for bidirectional medium flow.

**Figure 1.**
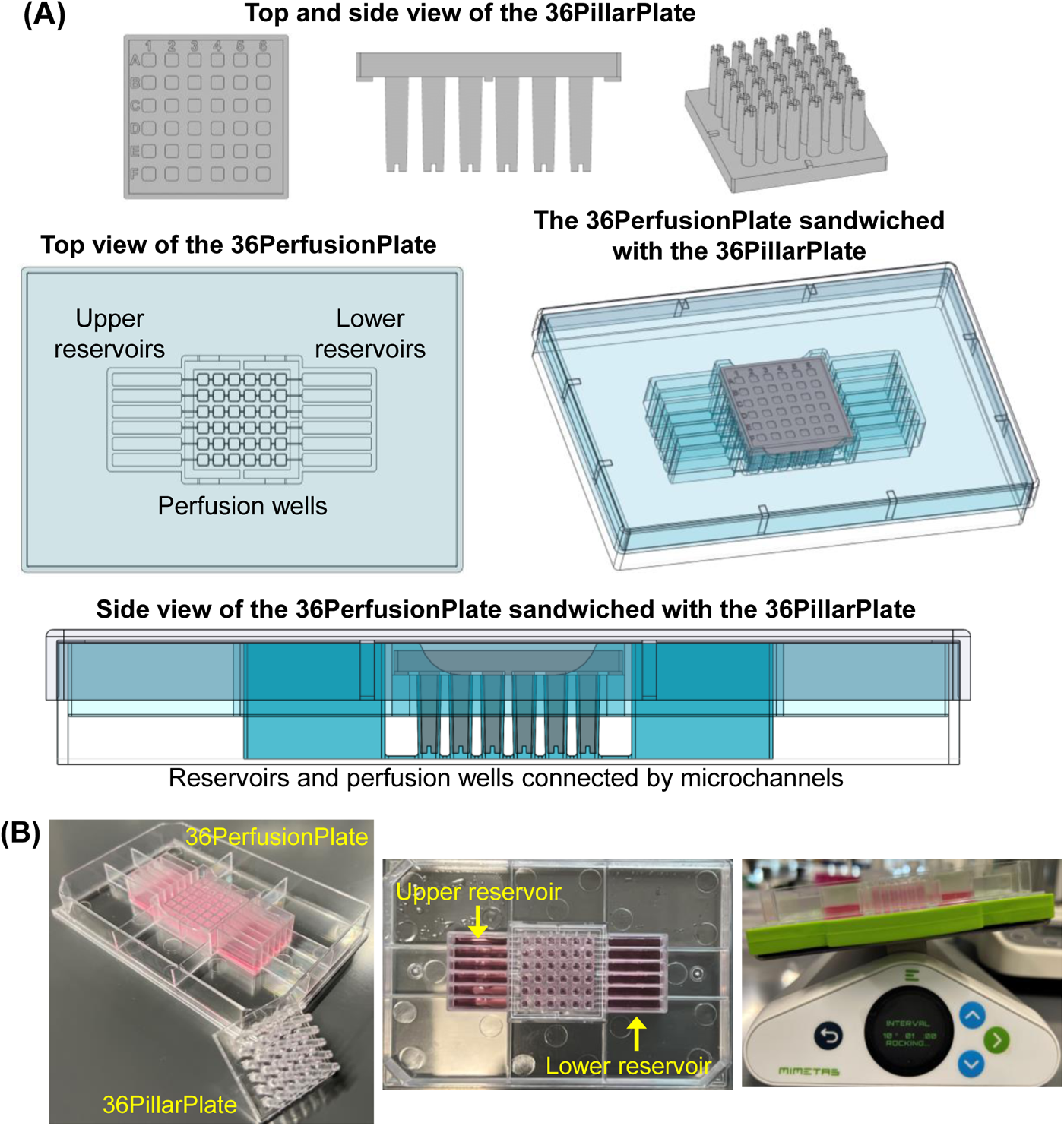
Design of a pillar/perfusion plate platform for dynamic organoid culture: **(A)** SolidWorks design of a 36PillarPlate with a 6 x 6 array of pillars, a 36PerfusionPlate with a 6 x 6 array of complementary perfusion wells, and the sandwiched pillar/perfusion plates. **(B)** Injection molded pillar/perfusion plates made of polystyrene and the use of a digital rocker to induce bidirectional fluid flow for dynamic cell culture.

### Human iPSC culture and maintenance

Human iPSC EDi029A (Cedar Sinai Biomanufacturing Center) was thawed, cultured, and passaged according to the manufacturer’s recommended protocol. Briefly, iPSCs with the initial passage number of 24 were cultured in a 6-well plate coated with 0.5 mg Matrigel™ (354230, Fisher Scientific) in complete mTeSR plus medium (100-0276, Stem Cell Technologies). The iPSCs were passaged every 4 - 5 days when it reached 80 - 90% confluency using the StemPro^®^ EZPassage™ disposable stem cell passaging tool (23181-010, Life Technologies). The cells were passaged as small clumps with a 1:6 split ratio and cultured with daily medium change. iPSCs at early passage numbers (up to 34 – 36) stored in a frozen vial were used for generating cerebral organoids.

### Formation of embryoid bodies (EBs) with iPSCs

iPSCs colonies were harvested using Accutase^®^ (A1110501, Gibco) and resuspended in complete human embryonic stem cell (hESC) medium consisting of DMEM/F12 (11320033, ThermoFisher), 20% (v/v) knockout serum replacement (10828010, ThermoFisher), 1% (v/v) GlutaMAX (35050-038, Invitrogen), 1% (v/v) MEM-nonessential amino acids (MEM-NEAA; M7145, MilliporeSigma), 3% hESC quality FBS (10439001, ThermoFisher), 100 µM 2-mercaptoethanol (21-985-023, Fisher Scientific), 4 ng/mL basic fibroblast growth factor (bFGF; 100-18B, Peprotech) and rho kinase inhibitor CEPT cocktail. The CEPT cocktail consists of 5 µM Emricasan (50-136-5235, Fisher Scientific), 50 nM Chroman 1 (7163, Tocris), 0.7 µM Trans-ISRIB (5284, Tocris), and polyamine supplement diluted at 1:1000 (P8483, MilliporeSigma). The collected cells were further dissociated into single cells through pipetting and seeded in an ultralow attachment (ULA) 384-well plate (ULA-384U-020, Nexcelom) at a seeding density of 1000 cells in each well containing 50 µL of the hESC medium. The cells were maintained in the hESC medium with CEPT until day 4 of culture, and later replaced with hESC medium without CEPT for an additional three days. The culture medium was changed every alternate day by removing half of the old medium and adding the same volume of the fresh medium. bFGF was added only at the first day of culture in hESC medium.

### Spheroid transfer to the 36PillarPlate from the ULA 384-well plate

On day 6, embryonic bodies (EBs) were formed in the ULA 384-well plate with the seeding density of 1000 cells/well using complete hESC medium. The EBs were subsequently transferred to the pillar plate with 5 µL of 6 - 8 mg/mL of Matrigel by using a simple sandwiching and inverting method ^36^. Briefly, the pillar plate with Matrigel was sandwiched onto the ULA 384-well plate with EBs. The sandwiched plates were then inverted and incubated for 30 - 40 mins at 37°C in a CO_2_ incubator to facilitate EB transfer and encapsulation in Matrigel.

### Differentiation and maturation of cerebral organoids on the 36PillarPlate in dynamic and static conditions

Cerebral organoids were differentiated on the 36PillarPlate by following the Lancaster protocol ^27^ with slight modification. Briefly, the EBs on the pillar plate were cultured in the neural induction medium (NIM) consisting of DMEM-F12, 1% (v/v) N2 supplement (17502001, ThermoFisher), 1% (v/v) GlutaMax, 1% (v/v) MEM-NEAA, and 1 µg/mL heparin (H3149, MilliporeSigma) in a 384DeepWellPlate (Bioprinting Laboratories Inc.) for 5 - 6 days with medium change every alternate day. After 5 - 6 days of EB culture in the NIM to generate neuroectoderm (NE), organoids were differentiated in cerebral organoid differentiation and maturation medium without vitamin A (CDM-vit A) either in the 384DeepWellPlate for static cerebral organoid culture or in the 36PerfusionPlate for dynamic organoid culture (**Figs. 2A & 2B**). The CDM-vit A consists of an equal volume of DMEM-F12 and neurobasal medium (21103049, Fisher Scientific) with 0.5% (v/v) N2 supplement, 1% (v/v) B27 supplement without vitamin A (12587010, ThermoFisher), 0.025% (v/v) insulin (I9278, MilliporeSigma), 1% (v/v) GlutaMax, 0.5% (v/v) MEM-NEAA, 1% (v/v) penicillin-streptomycin, and 50 µM 2-mercaptoethanol.

For dynamic cerebral organoid culture, 500 µL of CDM-vit A was added in both upper and lower reservoirs, corresponding to 1000 µL of the medium in each channel of the 36PerfusionPlate. The sandwiched pillar/perfusion plate containing cerebral organoids was placed on a digital rocker (MIMETAS) with 10° tilting angle and 1 minute frequency of angle change in the CO_2_ incubator at 37°C. The CDM-vit A medium was changed every alternate day till day 40 by tilting the sandwiched plate for 2 minutes, aspirating out the old medium from the lower reservoirs, and dispensing 500 µL of fresh CDM-vit A in both upper and lower reservoirs. On day 41, CDM-vit A was switched to cerebral organoid differentiation and maturation medium with vitamin A (CDM+vit A) with medium change every alternate day till day 63. The CDM+vit A consists of an equal volume of DMEM-F12 and neurobasal medium with 0.5% (v/v) N2 supplement, 1% (v/v) B27 supplement with vitamin A, insulin, 1% (v/v) GlutaMax, 0.5% (v/v) MEM-NEAA, 1% (v/v) penicillin-streptomycin, 100 µM 2-mercaptoethanol and 2% (v/v) of Matrigel. Previous studies have demonstrated that incorporating dissolved extracellular matrix, specifically Matrigel, in the culture resulted in distinct cortical plate formation while maintaining the integrity of the basement membrane ^37,38^.

For static organoid culture, 80 µL of CDM-vit A was added to each well of the 384DeepWellPlate. The pillar plate containing organoids was sandwiched onto the deep well plate and placed in the CO_2_ incubator at 37°C. The CDM-vit A medium was changed every alternate day till day 40. On day 41, CDM-vit A was switched to CDM+vit A with medium change every alternate day till day 63.

### Measurement of IC_50_ values with cerebral organoids and model compounds

Prior to assessing DNT potential of model compounds including ascorbic acid (DNT negative) and methylmercury (DNT positive), we determined IC_50_ values with cerebral organoids to determine acute toxicity of the model compounds. Briefly, the pillar plate with day 9 cerebral organoids was sandwiched onto the perfusion plate containing the compounds. The sandwiched plates were placed on the digital rocker at 10° tilting angle and 1 min frequency of angle change and incubated in the CO_2_ incubator at 37°C for 48 hours. The model compounds were 4-fold serially diluted in the NIM to prepare six different dosages. The concentration range used was 4 μM - 1 mM for ascorbic acid and 4 - 1024 nM for methylmercury, which was determined by using published C_max_ and IC_50_ values of the compounds ^12,15,39,40^. The six dosages of the compounds were dispensed in six upper reservoirs in the perfusion plate (one dosage per reservoir, six replicates of the perfusion wells per dosage). After 48 hours of compound exposure, organoid viability was measured by using a luminescence-based CellTiter-Glo 3D^®^ cell viability assay kit (G9681, Promega) and recording luminescence intensity with a microtiter well plate reader (BioTek® Cytation 5).

To plot dose-response curves and calculate IC_50_ values, the percentage of live organoids was calculated using the following equation:

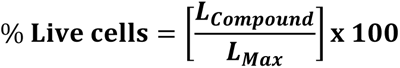

where L_Compound_ is the luminescence intensity of cerebral organoids exposed to a test compound and L_Max_ is the luminescence intensity of fully viable organoids (control). To produce a conventional sigmoidal dose-response curve with response values normalized to span the range from 0% to 100% plotted against the logarithm of test concentrations, we normalized the luminescence intensities of all cerebral organoid spots with the luminescence intensity of a 100% live organoid spot (cerebral organoids contacted with no compound) and converted the test compound concentration to their respective logarithms using Prism 9.3.1 (GraphPad Software, San Diego, CA, USA). The sigmoidal dose-response curve (variable slope) and IC_50_ value (i.e., concentration of test compound where 50% of organoid viability inhibited) were obtained using the following equation:

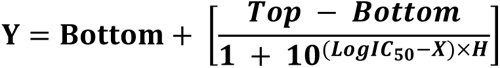

where IC_50_ is the midpoint of the curve, H is the hill slope, X is the logarithm of test concentration, and Y is the response (% live cells), starting from the top plateau (Top) of the sigmoidal curve to the bottom plateau (Bottom).

### Assessment of developmental neurotoxicity (DNT) potential with model compounds

For the assessment of DNT potential with the model compounds, day 9 cerebral organoids on the pillar plate were treated with 250 μM of ascorbic acid or 250 nM of methylmercury in the perfusion plate for 1 and 3 weeks. The test concentration of ascorbic acid and methylmercury for DNT potential was determined from the dose response curve (**Fig. 5B**), which was below their IC_50_ value for the assessment of long-term exposure. Briefly, the compounds dissolved in 1000 μL of the NIM were dispensed and equilibrated in each channel of the perfusion plate. The pillar plate with day 9 cerebral organoids was sandwiched onto the perfusion plate containing the compounds. The sandwiched plates were placed on the digital rocker at 10° tilting angle and 1 min frequency of angle change and incubated in the CO_2_ incubator at 37°C for 3 days. On day 3 of compound exposure, the compounds in the NIM were switched to the compounds in the CDM-vit A according to the Lancaster protocol of cerebral organoid differentiation. The compound exposure in the pillar/perfusion plate was continued for 1 and 3 weeks to study the effect of chronic exposure on organoid morphology and gene expression. The cerebral organoids cultured in the pillar/perfusion plate without compound were used as a negative control.

### Assessment of organoid viability

The viability of dynamically and statically cultured cerebral organoids on day 31 was measured with the CellTiter-Glo 3D^®^ cell viability assay kit as well as a fluorescence-based Live/Dead™ viability/cytotoxicity kit (L3224, Life Technologies) containing calcein AM and ethidium homodimer-1 (EthD-1) by following the manufacturers’ recommended protocol. For the CellTiter-Glo assay, the pillar plate with cerebral organoids was sandwiched onto an opaque white 384-well plate containing 50 µL/well of CellTiter-Glo^®^ reagent diluted with the culture medium at 3:1 ratio. The sandwiched plate was incubated for 1 hour at room temperature on an orbital shaker after which the pillar plate was discarded. Luminescence intensity from the opaque 384-well plate was measured using a microtiter well plate reader (BioTek^®^ Cytation 5).

In addition to the luminescence-based viability assay, a fluorescence-based cell viability assay was performed with calcein AM and EthD-1. Briefly, day 31 cerebral organoids on the pillar plate were rinsed with 1x PBS and stained with 80 μL of 2 μM calcein AM and 4 μM EthD-1 in DMEM/F-12 medium in the 384DeepWellPlate for 2 hours at 37°C in the CO_2_ incubator. After staining, the organoids were rinsed three times with PBS for 10 minutes each. The fluorescence images were obtained with a fully automated bright-field and fluorescence microscope (BZ-X800E, Keyence).

### Cryosectioning and hematoxylin and eosin (H & E) staining

The cerebral organoids were fixed with 4% paraformaldehyde (PFA) overnight at 4°C, washed with PBS for 10 minutes each for three times, and incubated with 30% sucrose overnight at 4°C. Cryoblocks of organoids were formed using Optimal Cutting Temperature Freezing Matrix (OCT, Sakura-Finetek, Torrence, CA) and were equilibrated to -18°C prior to cryosectioning. The frozen organoids were sectioned at 8 µm-thick slices on a Leica CM3050S cryostat (-18°C chamber, - 20°C object, 3.5° blade angle) and mounted to silanated microscope slides (Superfrost Plus, Fisher Scientific, Hampton, NH). The slides with sectioned organoids were dried under laminar flow at room temperature and kept at -80°C until staining.

The slides with sectioned organoids were individually thawed at room temperature under laminar flow for 30 minutes before staining. The slides were fixed in 10% buffered formalin for 20 minutes, washed with distilled water, and stained with Leica Selectech reagents (Hematoxylin 560, Alcoholic Eosin-Y 515, Richmond, IL). Stained slides were rinsed with distilled water and were dehydrated in alcohol. The slides were then cleared with xylene and covered with glass cover slips using permanent synthetic mounting medium. The images were obtained with a confocal microscope (LSM710, Zeiss) with 4x and 10x magnification.

### Immunofluorescence (IF) staining of whole cerebral organoids

All the solutions and reagents used for staining were dispensed at 80 µL/well in the 384DeepWellPlate. The pillar plate with cerebral organoids was sandwiched onto the deep well plate to perform the steps of cell staining. Briefly, dynamically, and statically cultured cerebral organoids on the pillar plate were rinsed with 1x PBS and treated with 4% PFA in the deep well plate overnight at 4°C. The organoids were permeabilized with 1% Triton X-100 in PBS for 1 hour at room temperature and then exposed to a blocking buffer (consisting of 4% normal donkey serum in 1x PBS with 0.5% Triton X-100) overnight at 4°C. Primary antibodies were diluted to recommended concentrations using the blocking buffer. The organoids were incubated with primary antibodies overnight at 4°C. After primary antibody staining, organoids were rinsed with 0.2% Triton X in PBS three times for 20 minutes each and incubated with appropriate secondary antibodies diluted in the blocking buffer overnight at 4°C. The stained organoids were washed with 0.5% Triton X-100 in PBS three times for 20 minutes each and incubated with 1 µg/mL DAPI in 0.5% Triton X-100 in PBS for 30 minutes. This was followed by final washing with PBS three times for 10 minutes each and incubation with a tissue clearing solution (Visikol Histo-M) for 1 hour. Images were obtained with a confocal microscope (LSM710, Zeiss) with 10x and 40x magnification and were analyzed with ImageJ/Fiji v1.54f software. The primary and secondary antibodies used are listed in **Supplementary Tables 1 and 2**, respectively.

### Calcium imaging of cerebral organoids

Organoids cultured for 63 days in static and dynamic conditions were stained with 10 µM of Fluo-4 AM cell permeant dye (F14201, ThermoFisher) for calcium imaging. Briefly, the culture medium containing 10 µM Fluo-4 AM dye was dispensed at 80 µL/well in the 384DeepWellPlate. The pillar plate with cerebral organoids was sandwiched onto the deep well plate and then incubated for 3 hours in the CO_2_ incubator at 37°C. Subsequently, the pillar plate was rinsed three times with the culture medium alone for 10 minutes each. Fluorescence images were obtained with the confocal microscope (LSM710, Zeiss) at 10x magnification.

### Gene expression analysis *via* qPCR

The cerebral organoids in Matrigel were isolated from the pillar plate using Cultrex organoid harvesting solution (3700-100-01, R&D Systems) according to the manufacturer’s recommended protocol, which allows non-enzymatic depolymerization of Matrigel. Briefly, the pillar plate with cerebral organoids were sandwiched onto the deep well plate containing 80 µL of Cultrex organoid harvesting solution. The sandwiched plates were incubated for 30 minutes at 4°C and then centrifuged at 100 rcf for 10 minutes to detach the organoids. Total RNA was isolated from iPSCs and organoids on days 16, 31, and 63 by using the RNeasy Plus Mini Kit (74134, Qiagen) with the manufacturer’s recommended protocol. cDNA was synthesized from 1 µg of RNA by following the protocol of cDNA conversion kit (4368814, Applied Biosystems). Real time PCR was performed using SYBR™ Green Master Mix (A25742, ThermoFisher) and forward/reverse primers from IDT Technology. The PCR cycle was run 25 - 35 times at 95°C denaturation for 30 sec, 58 - 62°C annealing for 45 sec, depending on primer pair, and 72°C extension for 30 sec by using QuantStudio™ 5 Real-Time PCR System (A28574, Applied Biosystems). The expression levels of targeted genes were normalized using the housekeeping gene glyceraldehyde 3-phosphate dehydrogenase (GAPDH). The primers used are listed in **Supplementary Table 3**.

### Flow cytometry analysis

Day 31 cerebral organoids cultured in a dynamic condition were enzymatically dissociated into single cells using Accutase for 10 minutes at 37°C in a CO_2_ incubator. The resulting single-cell suspension was collected in a 15 mL tube and fixed with 4% paraformaldehyde (PFA) for 10 minutes at room temperature. The cells were permeabilized with 0.1% Triton X-100 in PBS for 15 minutes, followed by blocking with 5% normal donkey serum (NDS) in 0.1% Triton X-100 in PBS for 30 minutes at room temperature. The cells were incubated with 5 µg/mL of primary antibodies against PAX6, FOXG1, TBR2, and CTIP2 for 1 hour at room temperature. Subsequently, the cells were washed with the blocking solution for three times, followed by incubation with 2.5 µg/mL of secondary antibodies such as donkey anti-rabbit IgG (H+L) highly cross-adsorbed secondary antibody, Alexa Fluor 488 and donkey anti-rat IgG (H+L) highly cross-adsorbed secondary antibody, Alexa Fluor 594 for 1 hour at room temperature. The cells were washed three times with PBS, and flow cytometry analysis was performed with Cytek Aurora spectral flow cytometer (Cytek^®^ Biosciences). The data were analyzed using FlowJo^TM^ v10 software (BD Biosciences).

**Figure 2.**
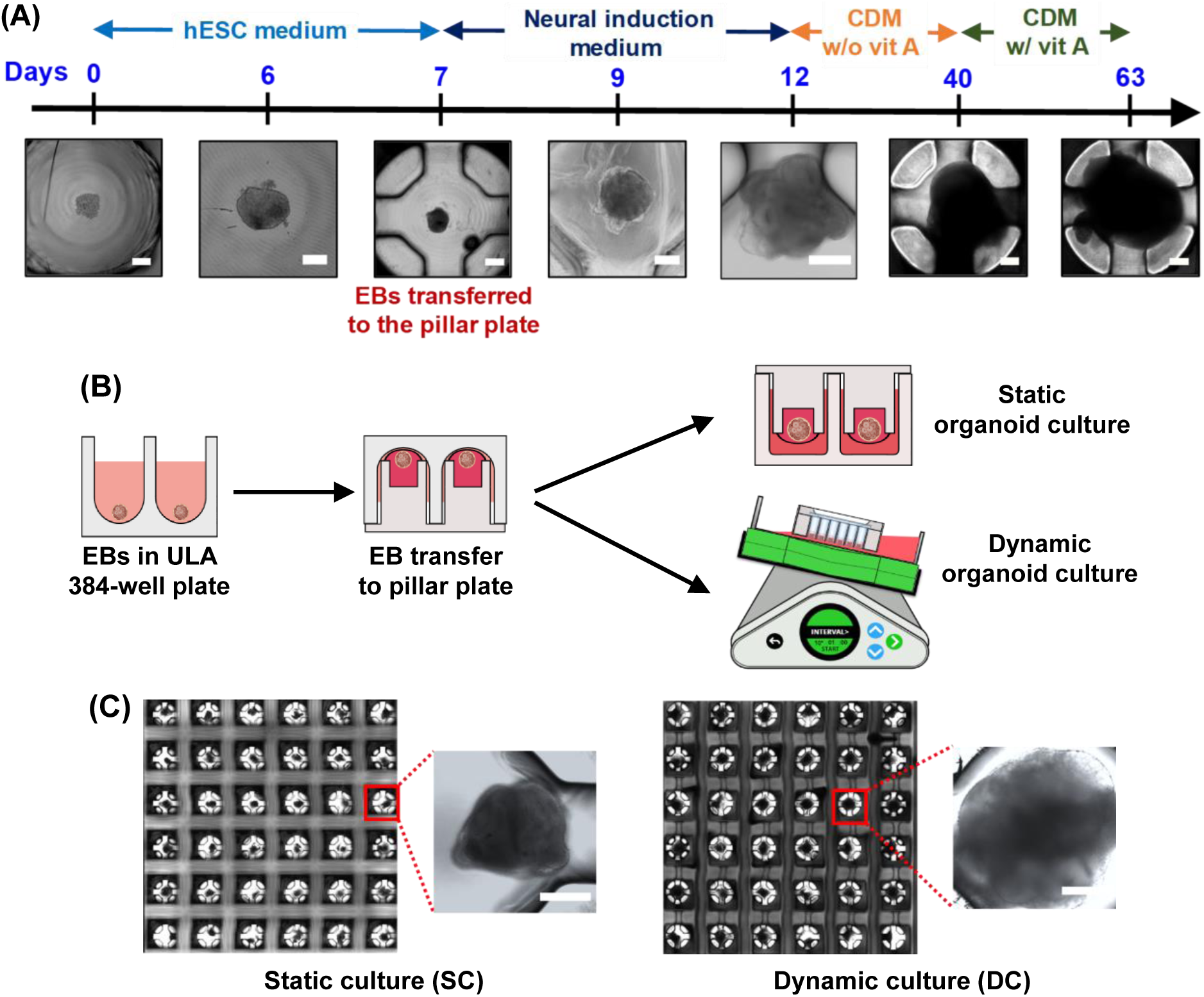
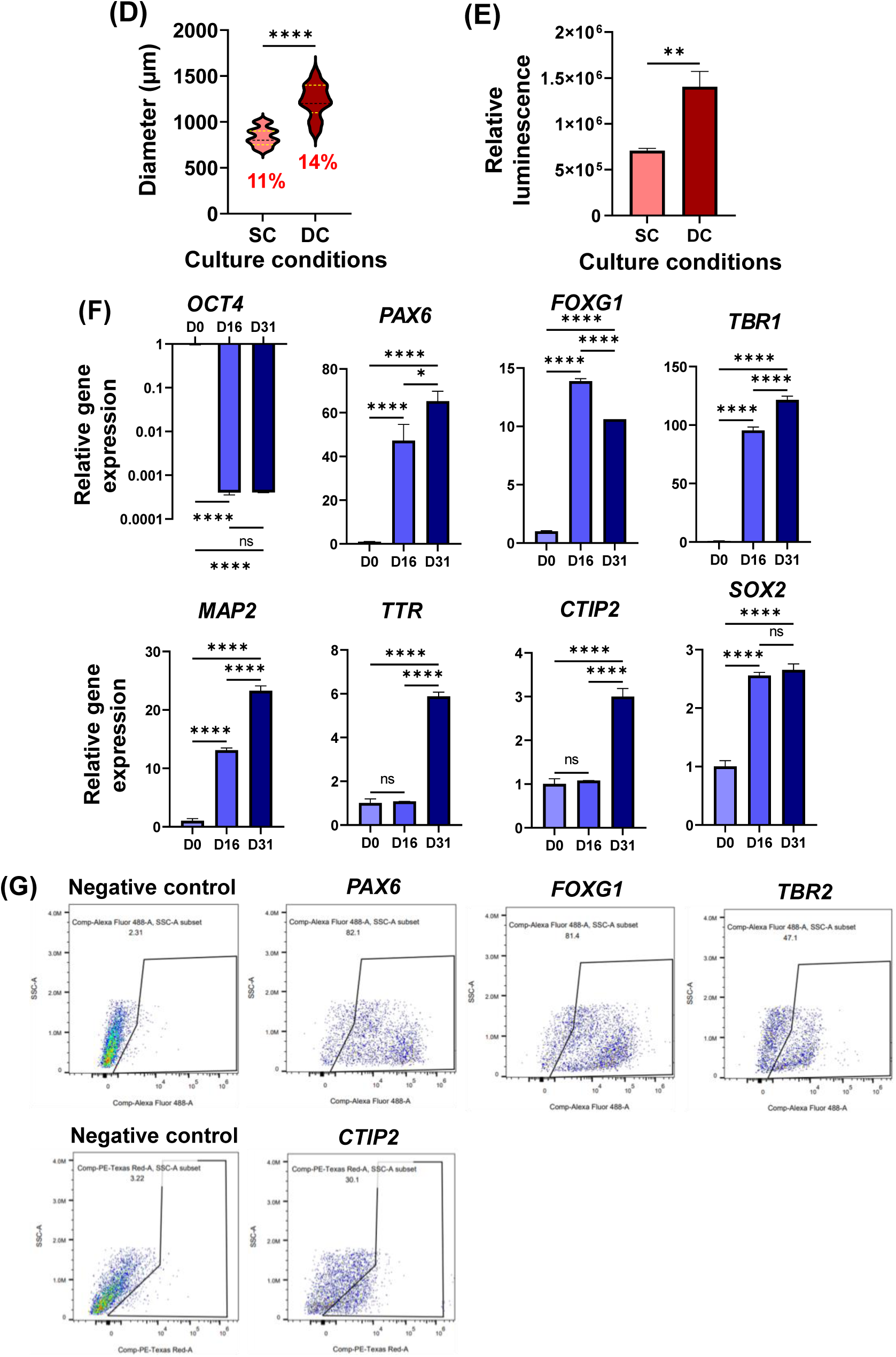
Dynamic culture of cerebral organoids in the pillar/perfusion plate: **(A)** The differentiation protocol of iPSCs into cerebral organoids. **(B)** Transfer of embryonic bodies (EBs) from an ultralow attachment (ULA) 384-well plate to a pillar plate as well as static and dynamic culture of cerebral organoids on the pillar plate. **(C)** Representative images of day 31 cerebral organoids cultured in a static condition using the 36PillarPlate/384DeepWellPlate and a dynamic condition using the 36PillarPlate/36PerfusionPlate. Scale bars: 350 μm. **(D)** The diameter of day 31 cerebral organoids cultured in static (SC) and dynamic (DC) conditions. n = 36 per condition. **(E)** ATP levels in day 31 cerebral organoids measured with CellTiter-Glo 3D cell viability assay kit. n = 18. **(F)** Neural biomarker expression in dynamically cultured cerebral organoids at days 0, 16, and 31, including *OCT4* pluripotency marker, *PAX6* forebrain neuroprogenitor marker, *FOXG1* forebrain marker, *TBR1* cortical preplate marker, *MAP2* mature neuronal marker, *TTR* choroid plexus marker, *CTIP2* deep cortical neuronal marker, and *SOX2* proliferating neuroprogenitor marker. Statistical significance was assessed by one-way ANOVA. **** for p < 0.0001, *** for p < 0.001, ** for p < 0.01, * for p < 0.05, n. s. = not significant (p > 0.05). n = 10 - 12 per qPCR run. **(G)** Flow cytometry analysis of day 31 cerebral organoids cultured in a dynamic condition to assess the percentage of the cells expressing PAX6, FOXG1, TBR2, and CTIP2. n = 8.

### Statistical analysis

Statistical analysis was performed using GraphPad Prism 9.3.1 (GraphPad Software, Inc., CA, USA). The data was expressed as mean ± SD. P values were calculated using one-way ANOVA analysis. The statistical significance threshold was set at **** for p < 0.0001, *** for p < 0.001, ** for p < 0.01, * for p < 0.05, and ns = not significant (p > 0.05). Sample sizes are indicated in the figure legends.

## Results

### The pillar/perfusion plate allowed simple and robust, dynamic cerebral organoid culture

To develop cerebral organoid models for assessing DNT potential of compounds, there was a need for generating organoids reproducibly and overcoming the issue of diffusion limitation. Here, we introduced the pillar/perfusion plate that supports the dynamic culture of cerebral organoids. Due to unique structure of the pillar with side walls and slits, it allowed robust transfer of EBs to the pillar plate ^36^. The single and uniform size of EBs transferred to the pillar plate allowed to generate cerebral organoids reproducibly with the coefficient of variation (CV) in the range of 7 - 19% in three trials ^36^. To demonstrate the cerebral organoid culture uniformly across the entire pillar plate, single EBs with a uniform size were transferred to each pillar while maintaining bidirectional flow under the pillars (**Fig. 2B**). The pillar/perfusion plate was designed to generate gravity-driven, ultrafast flow under the pillars by using Bernoulli’s principle, which improved the diffusion of nutrients into the organoids while operating it on a digital rocker in the CO_2_ incubator ^34,35^. The flow rate and shear stress generated in the pillar/perfusion plate were comparable to those in large capillaries or small veins ^34^. The average flow rate was 6.4 ± 0.3 µL/sec at 10° tilting angle, 1 minute frequency of angle change, and 1000 µL medium/channel ^34^. We generated cerebral organoids in the pillar/perfusion plate by using the Lancaster protocol ^27^ with slight modification (**Fig. 2A**). The culture of cerebral organoids was maintained in CDM-vit A until day 40. Subsequently, the medium was changed to CDM+vit A until day 63 (**Fig. 2A**). Dynamically cultured cerebral organoids in the pillar/perfusion plate were compared with their counterpart, statically cultured cerebral organoids in the pillar/deep well plate to demonstrate the effect of rapid fluidic flow on organoid maturity (**Fig. 2B**). Interestingly, the size of dynamically cultured cerebral organoids on day 31 was in the range of 1000 - 1400 µm in diameter, which was significantly bigger than the size of statically cultured cerebral organoids in the range of 700 - 900 µm with the CV value less than 15% (**Figs. 2C & 2D**). The size of the organoids after 63 days of static and dynamic cultures was approximately 1200 - 1500 µm and 2000 - 2500 µm in diameter, respectively. The ATP content measured using CellTiter-Glo assay in the day 31 organoids cultured in the dynamic condition showed a greater than 2-fold increase compared to their static counterpart, indicating higher number of cells in dynamic culture (**Fig. 2E**). In addition, there were only a few dead cells found in the day 31 cerebral organoids cultured in the static and dynamic conditions, which were confirmed by cell staining with calcein AM and EthD-1 (**Supplementary Fig. 1**). Furthermore, cryosectioning of day 31 cerebral organoids and H & E staining revealed dense cellular organization in the dynamically cultured organoids as compared to the statically cultured organoids, indicating enhanced cell proliferation in dynamic culture (**Supplementary Fig. 2A**). The cryo-sectioned cerebral organoids cultured for 31 days in a dynamic condition revealed the presence of distinct layers within the organoids, which was confirmed by the spatial separation between the mature neuronal marker *TUBB3* and the proliferating neuroprogenitor marker *SOX2* (**Supplementary Fig. 2B**). Finally, qPCR data analysis showed increased organoid maturity along with the culture period in the dynamic condition (**Fig. 2F**). A 1000-fold decrease in the expression of the pluripotency marker *OCT4*, along with a 60-fold increase in the expression of the neuroprogenitor marker *PAX6* indicated the successful neural induction in cerebral organoids. *PAX6* expression is important for initiating neurogenesis along with proliferation of neural stem cell ^41^. The increased expression of *PAX6* by more than 40-fold on day 16 showed efficient neural commitment within the organoids. In addition, a 10-fold increase in the forebrain marker *FOXG1*, a 120-fold increase in the cortical preplate marker *TBR1*, a 25-fold increase in the mature neuronal marker *MAP2*, a 6-fold increase in the choroid plexus marker *TTR*, and a 3-fold increase in the deep cortical neuronal marker *CTIP2* in day 31 cerebral organoids indicated the successful neuronal differentiation, brain regionalization, and cortex identities in dynamic culture. These results highlight that the pillar/perfusion plate can enhance organoid growth in dynamic culture and organoid maturity with longer differentiation time. In addition, the flow cytometry analysis of day 31 cerebral organoids cultured in a dynamic condition revealed different types of cellular expression within the organoids, having 82.1% PAX6+, 81.4% FOXG1+, 47.1% TBR2+, and 30.1% CTIP2+, respectively (**Fig. 2G**).

### Dynamic culture enhanced differentiation and maturation of cerebral organoids

Here, we conducted tests to determine if dynamic culture in the pillar/perfusion plate could enhance organoid maturity. Briefly, cerebral organoids were differentiated in static and dynamic conditions for 31 and 63 days and characterized by qPCR and immunofluorescence staining (**Figs. 3 and 4**). Overall, qPCR data analysis of day 31 and day 63 organoids revealed a significant enhancement in organoid maturity along with the time of culture in dynamic culture compared to static culture (**Fig. 3**). The expression of *PAX6* forebrain neuroprogenitor marker, *TBR2* intermediate progenitor marker, *FOXG1* forebrain marker, and *CTIP2* cortical neuronal marker were significantly increased in dynamically cultured organoids for 31 days as compared to their static counterpart. In addition to *PAX6*, *TBR2*, and *FOXG1*, the expression levels of *TTR* choroid plexus marker*, TBR1* cortical preplate marker, *CTIP1* postmitotic neuronal marker, *CTIP2* cortical neuronal marker*, TUBB3* mature neuronal marker*, SYN1* synaptic marker, *GFAP* astrocyte marker, *DCX* neuronal migration marker, *VGLUT1* excitatory neuronal marker, *MAP2* mature neuronal marker*, GABRG3* encoding a gamma-aminobutyric acid (GABA) receptor, and *SATB2* upper layer neuronal marker in the organoids were significantly enhanced in day 63 cerebral organoids in dynamic culture compared to those in static culture. The increased expression of *PAX6* along with the time of differentiation suggests that the organoids are becoming more mature and attaining the cerebral cortex specification ^41,42^. *PAX6* plays an essential role in the development of cerebral cortex and its increased expression implicates that the organoids are developing more specialized cell types ^42^. The intermediate progenitor marker *TBR2* is essential for the cortical neurogenesis expansion ^43,44^, and thus the increased expression of *TBR2* along with the time of differentiation indicates the initiation of cortical neurogenesis. The decreased expression of *FOXG1* from day 31 to day 63 could indicate the initiation of neural differentiation, as its expression is reported to fluctuate throughout cortical development and transiently decrease when neuronal differentiation is initiated from progenitors ^45^. The increment in the expression levels of *TTR*, *TBR1*, *CTIP1*, *CTIP2, TUBB3*, *SYN1*, *GFAP*, *DCX*, *VGLUT1*, *MAP2*, *GABRG3,* and *SATB2* in dynamic culture in comparison to the static counterpart demonstrated that the organoid was becoming more matured in dynamic culture. Moreover, the expression levels of these crucial markers were higher in day 63 cerebral organoids compared to day 31 cerebral organoids in both static and dynamic culture conditions, indicating the enhancement in organoid maturity along with the time of differentiation.

**Figure 3.**
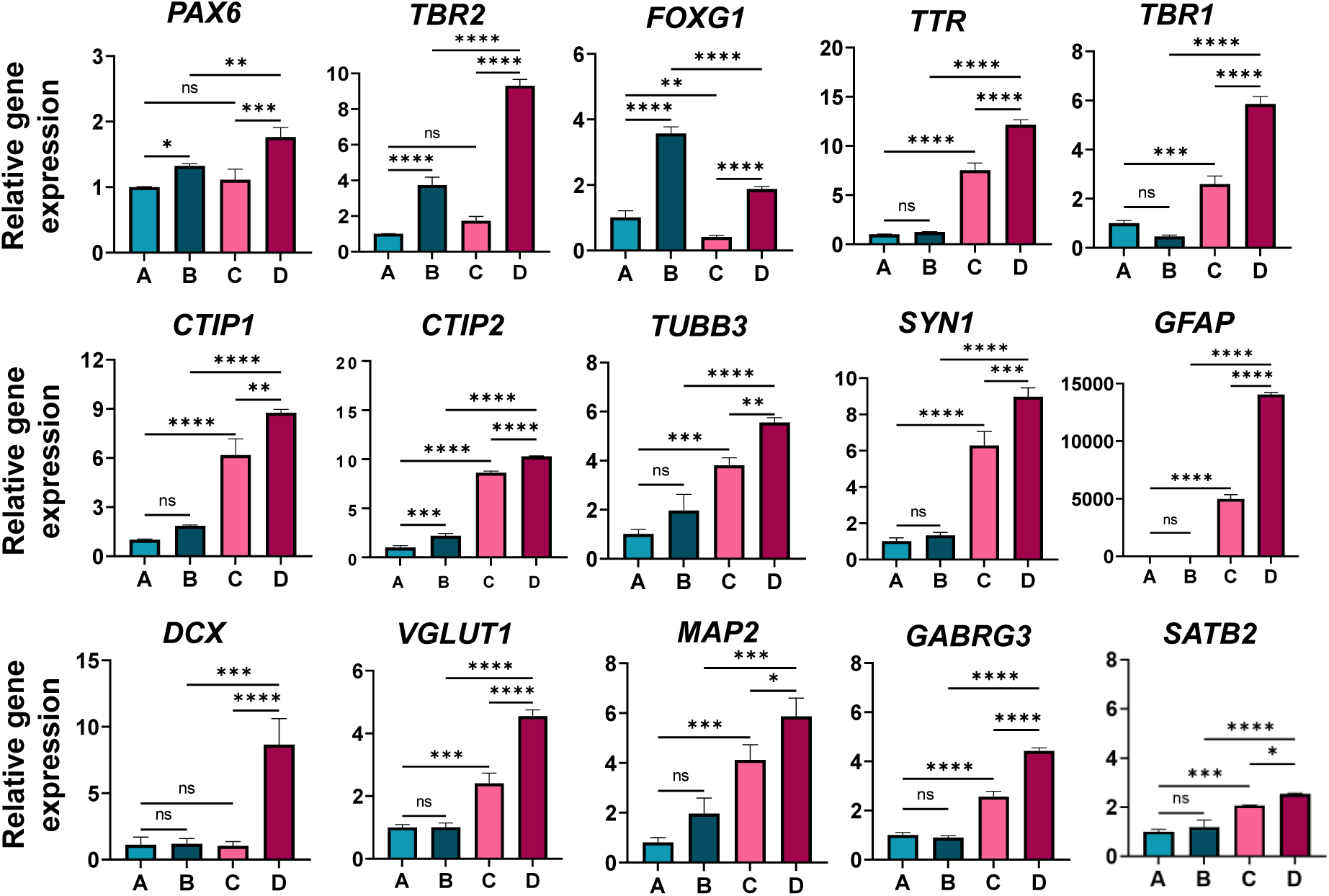
Relative neural biomarker expression in cerebral organoids in the pillar/perfusion plate: **(A)** statically cultured cerebral organoids for 31 days, **(B)** dynamically cultured cerebral organoids for 31 days, **(C)** statically cultured cerebral organoids for 63 days, **(D)** dynamically cultured cerebral organoids for 63 days. Neural biomarkers include *PAX6* forebrain neuroprogenitor marker, *TBR2* intermediate progenitor marker, *FOXG1* forebrain marker, *TTR* choroid plexus marker*, TBR1* cortical preplate marker*, CTIP1* postmitotic neuronal marker, *CTIP2* cortical neuronal marker*, TUBB3* mature neuronal marker*, SYN1* synaptic marker, *GFAP* astrocyte marker, *DCX* neuronal migration marker*, VGLUT1* excitatory neuronal marker, *MAP2* mature neuronal marker*, GABRG3* encoding a GABA receptor, and *SATB2* upper layer neuronal marker. Statistical significance was performed by one-way ANOVA. **** for p < 0.0001, *** for p < 0.001, ** for p < 0.01, * for p < 0.05, n. s. = not significant (p > 0.05). n = 10 - 12 per qPCR run.

In addition to qPCR analysis, immunofluorescence staining of whole cerebral organoids cultured in static and dynamic conditions for days 31 and 63 was performed to assess the microarchitecture of the cerebral organoids (**Fig. 4**). The stained organoids cultured in static and dynamic conditions displayed the higher expression of several forebrain region biomarkers, including PAX6, SOX2, CTIP2, TBR2, MAP2, and GFAP in dynamic culture compared to static culture, further illustrating the enhancement of organoid differentiation and maturation in dynamic culture. Overall, much clear cortical organization was observed in dynamically cultured organoids as CTIP2 and SOX2 staining revealed the distinctive arrangement of cortical neurons in the cortical plate-like region from the ventricular zone (**Fig. 4B**). It is well known that SOX2+ cells are present in the ventricular zone whereas CTIP2+ cells are in the deep cortical plate ^46^. Additionally, TBR2 staining revealed that the precursor cell of the cortical plate was dispersed at the periphery of the ventricular zone (**Fig. 4C**). MAP2 staining showed increased emergence of the basal neural layer in dynamic culture as compared to static culture (**Fig. 4D**). Moreover, GFAP staining of day 63 organoids showed higher expression of the astrocyte marker in dynamic culture (**Fig. 4E**). The astrocyte marker GFAP, mostly expressed at a late stage of differentiation ^37^, was observed after 2 months of differentiation on the pillar plate. The quantitative expression of the fluorescence intensity was also higher in dynamically cultured cerebral organoids in comparison to their static culture counterparts (**Supplementary Fig. 3**). The fluorescence intensity of each biomarker was normalized with the fluorescence intensity of DAPI. Furthermore, increased fluorescence was observed in neurons from calcium imaging in dynamic culture which is an indicator of increased intracellular calcium ion (**Supplementary Fig. 4**). Since calcium signal is essential for neurotransmitter release and synaptic plasticity, this result indicates that neurons within the cerebral organoids cultured in the dynamic condition have become more mature in comparison to their static counterpart.

**Figure 4.**
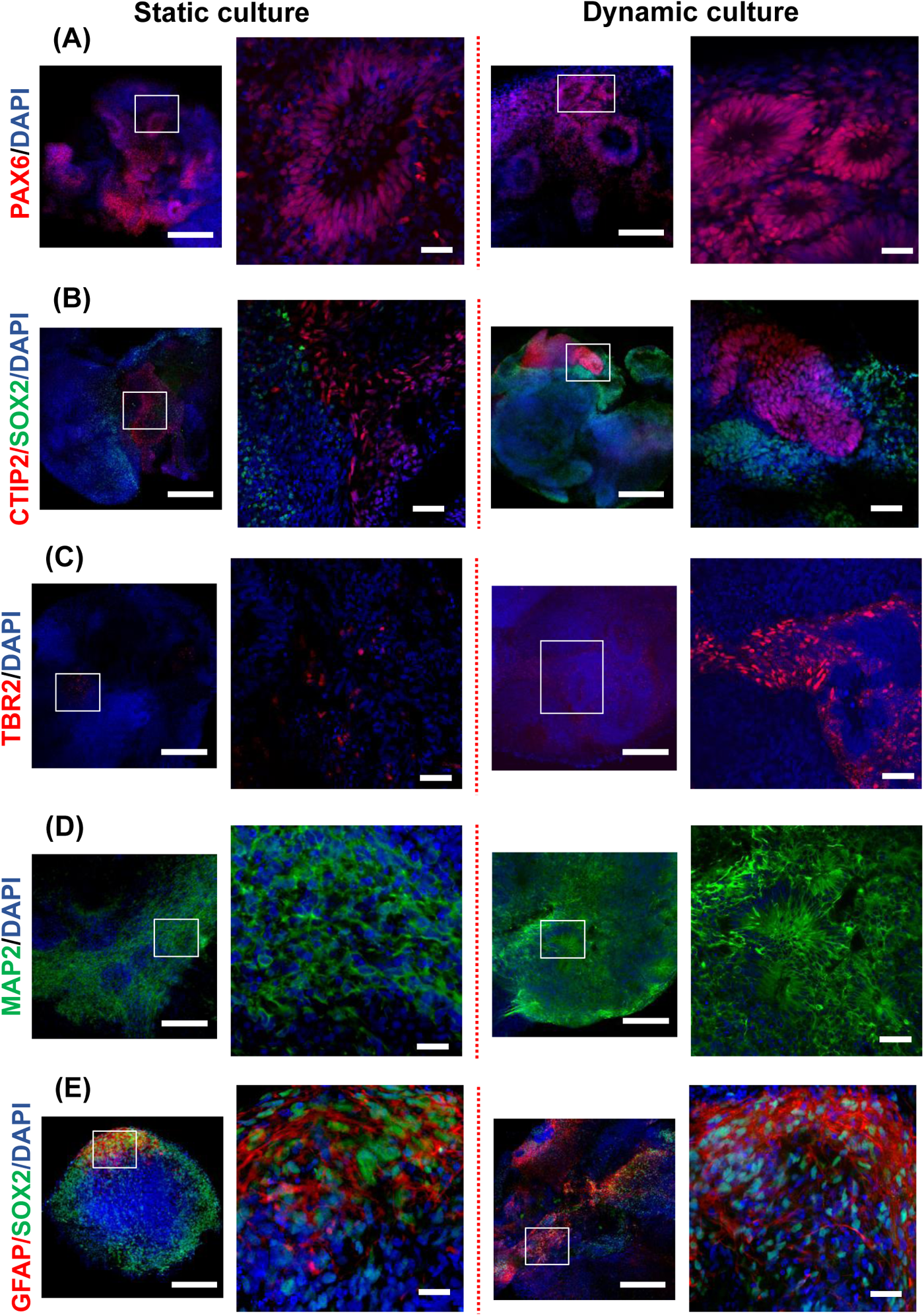
Immunofluorescence staining of cerebral organoids cultured in static (left panel) and dynamic (right panel) conditions: **(A)** PAX6, **(B)** CTIP2/SOX2, **(C)** TBR2, **(D)** MAP2, and **(E)** GFAP/SOX2. (A), (B), (C), and (D) are from day 31 cerebral organoids and (E) is from day 63 cerebral organoids. Neural biomarkers include PAX6 forebrain neuroprogenitor marker, CTIP2 deep cortical neuronal marker, SOX2 proliferating neuroprogenitor marker, TBR2 intermediate progenitor marker, MAP2 mature neuronal marker, and GFAP astrocyte marker. Scale bars: 200 µm and 50 µm (magnified).

### Assessment of developmental neurotoxicity (DNT) with dynamically cultured cerebral organoids in the pillar/perfusion plate

To prevent cell death from acute cytotoxic effects and accurately assess the potential for DNT following long-term compound exposure, it was necessary to measure acute toxicity of the model compounds and use dosages below their IC_50_ value. Briefly, day 9 cerebral organoids were treated with an appropriate concentration range of the model compounds, including ascorbic acid (DNT negative) and methylmercury (DNT positive), in the dynamic condition for 48 hours to determine the dose-response curves and calculate the IC_50_ values (**Fig. 5**). The average IC_50_ values obtained from two trials were 350 nM for methylmercury whereas ascorbic acid did not show cytotoxic effects at the tested concentration range.

**Figure 5.**
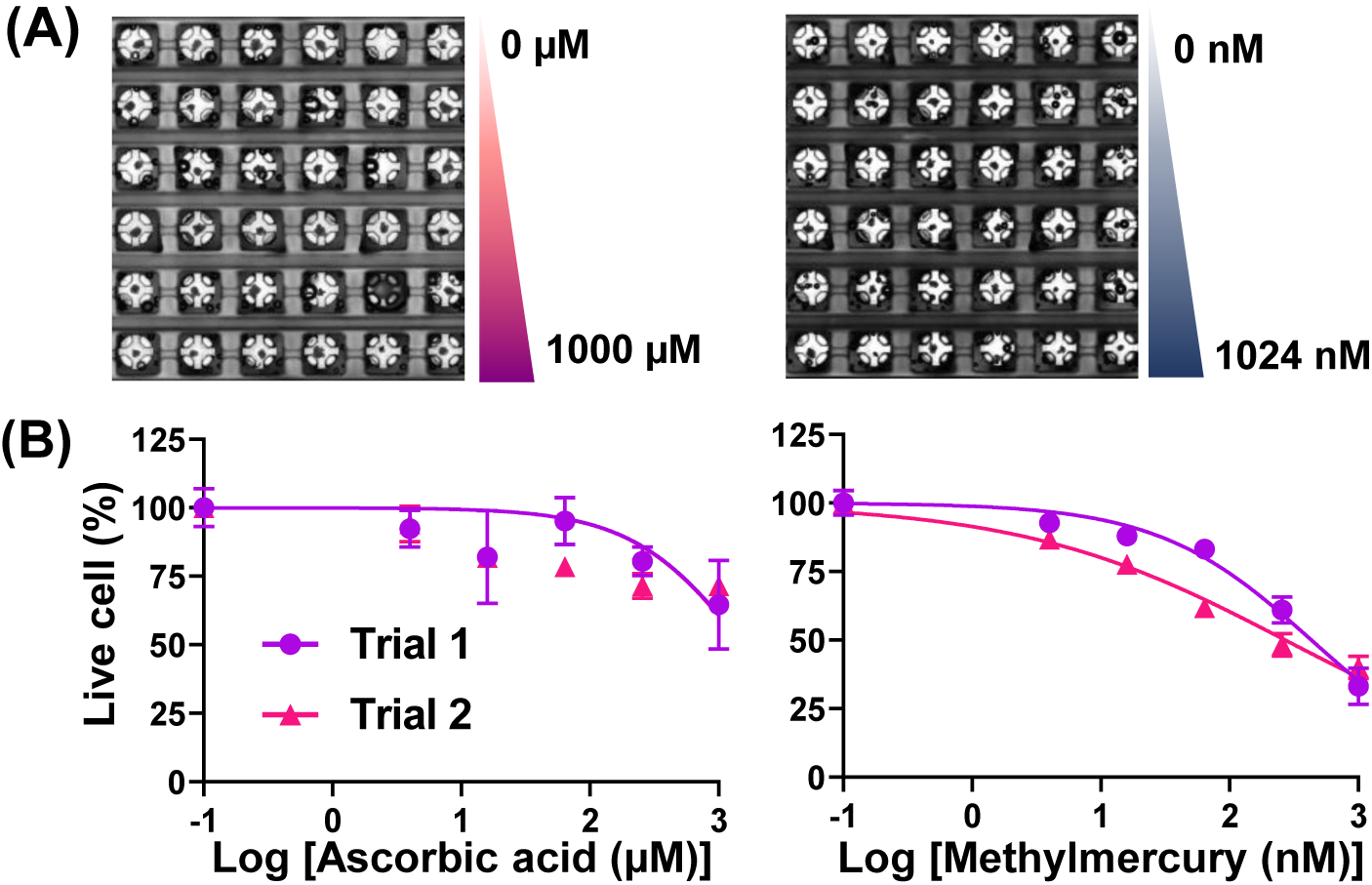
Acute toxicity of model compounds tested with cerebral organoids: **(A)** Stitched images of compound-treated cerebral organoids on the pillar plate for 2 days. **(B)** Dose-response curves of the model compounds tested. The IC_50_ values of methylmercury from two trials were 350 nM whereas ascorbic acid was noncytotoxic at the concentrations tested. n = 6 per dosage.

To investigate the DNT potential and the effects of chronic exposure to the model compounds, we selected the concentration below their IC_50_ value. The changes in organoid morphology and neural gene expression were assessed after 1 and 3 weeks of compound exposure in dynamic culture (**Fig. 6A**). Briefly, day 9 cerebral organoids in the pillar/perfusion plate were exposed to 250 µM ascorbic acid and 250 nM methylmercury for 1 and 3 weeks in the dynamic condition with medium change every alternate day. In addition to morphology change, the expression levels of representative neural biomarkers, including *PAX6* forebrain neuroprogenitor, *SOX2* proliferating neuroprogenitor, *FOXG1* forebrain, *NESTIN* neuronal progenitor cell, *TUBB3* neuronal cytoplasm, and *MAP2* mature neuron as well as *HMOX1* cellular stress marker were measured after 1 and 3 weeks of compound exposure to study the effect of long-term chemical exposure in the developing cerebral organoids. As anticipated, the exposure of day 9 cerebral organoids to 250 µM ascorbic acid for 1 and 3 weeks in the dynamic condition did not result in any significant changes in organoid morphology as well as neural gene expression as compared to the no compound exposure control (**Fig. 6B**). This result indicates that ascorbic acid doesn’t induce DNT. On the other hand, the exposure of day 9 cerebral organoids to 250 nM methylmercury for 1 and 3 weeks in the dynamic condition resulted in significant changes in organoid morphology and neural gene expression (**Fig. 6C**), indicating that methylmercury does induce DNT. The organoid size significantly decreased after 1 and 3 weeks of methylmercury treatment compared to the no compound exposure control. This could be due to the adverse effect of methylmercury on DNA methylation, resulting in impaired cell programming during long-term exposure ^47^. Additionally, the expression level of *FOXG1* forebrain marker increased 5-fold and 3-fold after 1 and 3 weeks of methylmercury exposure, respectively, compared to the no compound exposure control. Furthermore, the expression level of *HMOX1* cellular stress marker increased 3-fold after 3 weeks of methylmercury exposure. It was reported that the overexpression of *FOXG1* gene could be one of the causative factors of autism spectrum disorder ^48^. Therefore, our results suggest that early exposure to methylmercury in brain development may induce DNT. Additional qPCR analysis revealed that methylmercury exposure could lead to the decreased expression of *BCL2* anti-apoptotic marker and the increased expression of pro-apoptotic markers including *BAX*, *Caspase3*, and *Caspase9* (**Fig. 6D**). This result suggests that methylmercury exposure could induce apoptosis in long-term dynamic culture. Interestingly, day 9 cerebral organoids exposed to 250 nM methylmercury for 3 weeks in the static condition in the pillar/deep well plate did not induce any significant changes in organoid morphology (**Supplementary Fig. 5**). This could be attributed to the diffusion limitation of methylmercury in the core of the organoids in the static condition, highlighting the importance of using a dynamic culture system for compound exposure in DNT assessment.

**Figure 6.**
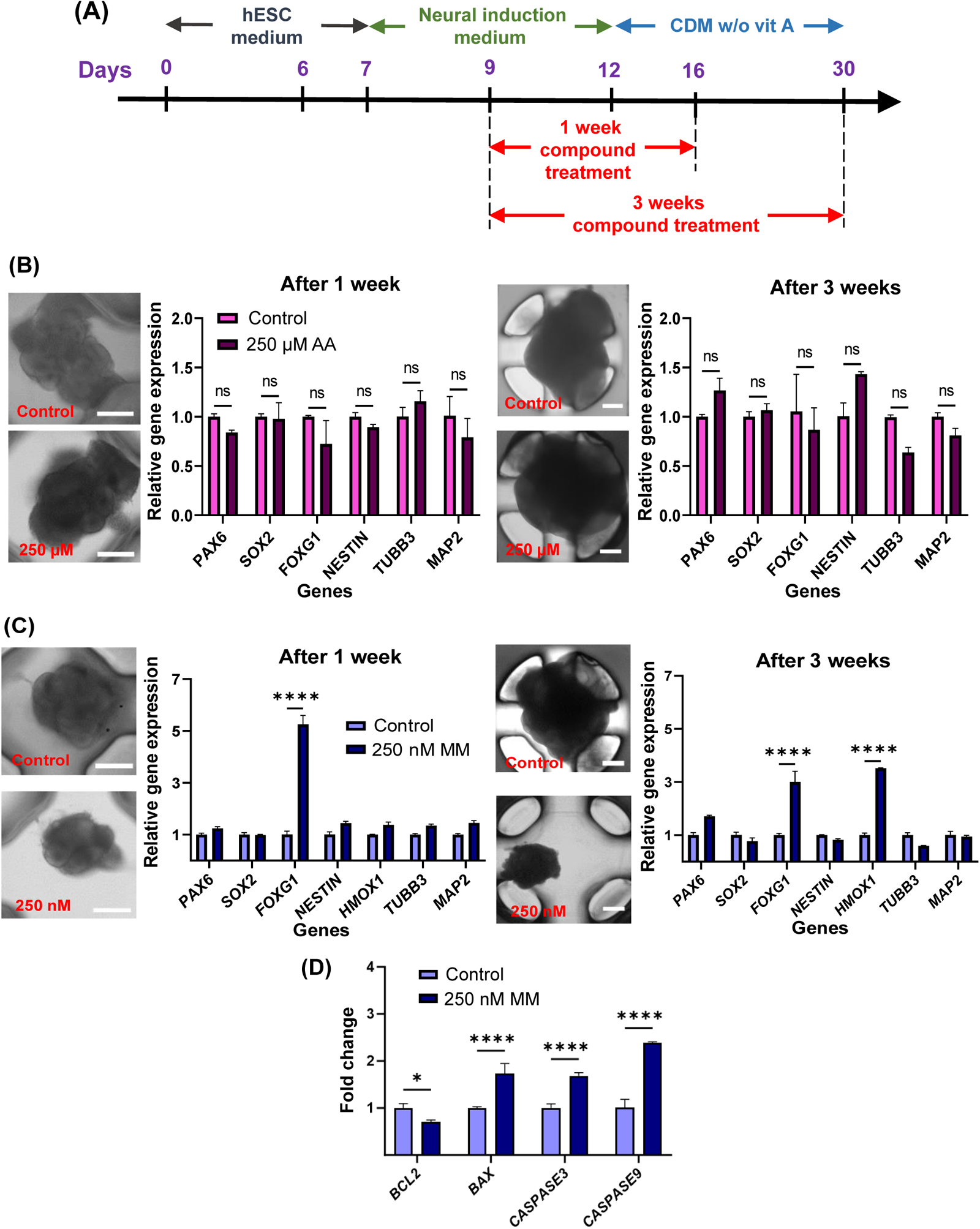
Developmental neurotoxicity of model compounds tested for 1 and 3 weeks with cerebral organoids in a dynamic condition: **(A)** The timeline of compound treatment with cerebral organoids. **(B)** Treatment of 250 µM ascorbic acid, and **(C)** Treatment of 250 nM methylmercury. Neural biomarkers measured after 1 and 3 weeks of compound exposure include *PAX6* forebrain neuroprogenitor marker, *SOX2* proliferating neuroprogenitor marker, *FOXG1* forebrain marker, *NESTIN* neuronal progenitor cell marker, *HMOX1* cellular stress marker, *TUBB3* neuronal cytoplasm marker, and *MAP2* mature neuronal marker. Scale bars: 200 µm. n = 12. **(D)** qPCR analysis of methylmercury-treated cerebral organoids for 3 weeks and no compound-treated organoids (control) for apoptotic biomarkers. n = 10 - 12 per qPCR run. *BCL2* represents an anti-apoptotic marker whereas *BAX*, *Caspase3*, and *Caspase9* represent pro-apoptotic markers.

## Discussion

Cerebral organoids generated from iPSCs have potential to model brain disorder and assess DNT of test compounds *in vitro* as they could mimic the early phase of brain development. Previous studies have demonstrated that perfused cell cultures enhance the maturity and complexity of organoids derived from hPSCs/hiPSCs ^29,30,49^. Additionally, several studies have demonstrated improved reprogramming and differentiation of various cell types in dynamic culture, addressing the formation of the necrotic core in long-term cultured, large organoids by improving oxygen and nutrient diffusion ^30,33,50–52^. Conventional methods of dynamic organoid culture typically rely on 6-well plates and petri dishes on an orbital shaker and spinner flasks, which require substantial resources in terms of cells and culture media and are low throughput for organoid imaging and compound testing. Recently, organoids have been combined with microphysiological systems to closely mimic the complexity of human tissues ^53^. Cho *et al.* demonstrated that dynamic culture in a pump-free microfluidic device improved nutrient and oxygen diffusion to the core, thereby increased the viability of brain organoids ^30^. They also demonstrated that bidirectional fluid flow enhanced the expansion of radial glial cells with neuroprogenitor cell (NPC) proliferation in an early stage of brain organogenesis ^30^. Although organoids can be cultured in microfluidic devices with minimal resources, they often require complex arrangements of pumps and tubes, making them user unfriendly and low throughput. Additional limitations, including inconvenient medium change, difficulty in cell staining and imaging, relatively high variation in organoid function, poor reproducibility, and non-specific adsorption of test compounds to polydimethylsiloxane (PDMS) used for chip fabrication, hinder the broader adoption of microfluidic devices in organoid culture for compound screening ^54^.

To address these limitations, we introduced the pillar/perfusion plate and demonstrated dynamic culture of cerebral organoids using gravity-driven, bidirectional flow on a digital rocker to enhance organoid maturity. With the sandwiching and inverting method developed for spheroid transfer, the 36PillarPlate and the 36PerfusionPlate supported dynamic culture of a 6 x 6 array of cerebral organoids per pillar plate. This platform miniaturizes dynamic organoid culture, requiring only 3.6 – 6 mL of cell culture media/perfusion plate every alternate day to generate 36 organoids (100 - 167 µL/organoid) whereas organoids in the 24-well plate and the spinner flask require 12 – 24 mL of cell culture media/well plate (500 – 1000 µL/organoid) every alternate day and 75 – 100 mL of culture media/flask every week, respectively. Thus, there is at least 5-fold up to 100-fold reduction in organoid culture. In addition, there is no need to remove organoids from the pillar plate for cell staining and imaging, which is critical for organoid-based compound screening. The pillar/perfusion plate built on the footprint of 384-well plates is compatible with existing lab equipment such as microtiter well plate readers and confocal microscopes for functional analysis of organoids. Furthermore, there is no concern for non-specific adsorption of compounds in the pillar/perfusion plate because it is made by injection molding with polystyrene.

Dynamically cultured cerebral organoids in the pillar/perfusion plate for 1 – 2 months exhibited higher proliferation and maturity compared to their static counterpart cultured in the pillar/deep well plate. We have already compared the expression levels of neuronal marker genes in statically cultured cerebral organoids in the pillar/deep well plate with those cultured in conventional Matrigel domes in the 24-well plate and demonstrated improved organoid maturity in the pillar/deep well plate due to culture miniaturization in our previous study ^36^. In the current work, dynamically cultured cerebral organoids showed increased expression levels of several genes including *PAX6*, *TBR2*, *FOXG1*, *TTR*, *TBR1, CTIP1*, *CTIP2*, *TUBB3*, *SYN1*, *GFAP*, *DCX*, *VGLUT1*, *MAP2*, *GABRG3,* and *SATB2*. In addition, dynamic organoid culture in the pillar/perfusion plate enhanced cell proliferation and viability as compared to static organoid culture. The improvement in viability and maturity of dynamically cultured organoids could be due to the enhanced diffusion of nutrients and oxygen and shear stress imposed on organoids *via* the bidirectional medium flow. We have previously demonstrated the improved cell proliferation and viability of Hep3B spheroids cultured in the pillar/perfusion plate due to improved diffusion of nutrients into the core of the spheroids ^34^. It is known that the diffusion rate of fluorescein increases significantly over time in perfused culture conditions, which was tested with a sodium fluorescein tracer across the gel ^55^. In addition, the role of mechanical and physical cues in organogenesis during early development has been well-established ^56^. Studies have reported the improved cell differentiation under the exposure of shear stress ^57^.

Although cerebral organoids have been used recently to model neurodevelopmental disorders such as microcephaly, Zika virus infection, and autism ^27,46,48^, very few studies have focused on studying the DNT potential of chemicals ^58,59^. Among the published studies related to the assessment of DNT, most of them have focused primarily on screening acute toxicity of chemicals rather than examining the adverse effects of chronic exposure to compounds on the developing brain. For example, 3D neurospheres and hESC-derived 3D neural tissue models have been used to study the adverse effects of short-term exposure to methylmercury ^11,12^. Human NPC-derived neurospheres were exposed to methylmercury for 48 hours, which resulted in decreased neuronal migration with fewer neuronal-like cells ^12^. In addition, hESCs-derived neurospheres were used to assess neurotoxicity through compound treatment for 72 hours ^15^. Furthermore, 2-month-old cerebral organoids were exposed to different concentrations of ethanol for 6 hours to simulate the adverse effects of excessive drinking by measuring signaling pathways and neural pathology phenotypes ^59^. Finally, day 10 cerebral organoids were exposed to ethanol for 14 days to study alcohol-induced pathogenesis ^58^, which revealed that ethanol could alter neurogenesis and cause an imbalance between excitatory and inhibitory neurons with hyper differentiation of glutamatergic neurons. Microfluidic devices have also been employed to assess acute neuronal toxicity by chemicals ^60,61^. Despite the extensive efforts to generate 3D cell/tissue models of the brain, it is still challenging to assess DNT potential of chemicals in a high-throughput screening (HTS) platform, particularly in a dynamic culture system, due to limitations of conventional organoid culture systems. Recently, Renner *et al.* demonstrated high-throughput compound screening with midbrain organoids in an ultralow attachment (ULA) 96-well plate to evaluate general neurotoxicity and dopaminergic neuron-specific toxicity with 84 compounds ^16^. Nonetheless, midbrain organoids were derived from neural precursor cells in scaffold-free culture, relying on patterning molecules for guided differentiation, which could not be generally applicable to other organoids that require hydrogel encapsulation to promote self-organization and differentiation of cultured stem cells ^62^. Thus, most of *in vitro* DNT assessments typically involve short-term compound exposure where cellular responses are measured at a single time point for acute dose responses. This approach often overlooks the potential of long-term cumulative effects. To enhance the broader utility of *in vitro* 3D cell/tissue models in risk assessment, the cell culture systems should be able to identify and characterize the long-term, adverse effects of chemicals, arising from repetitive exposure over extended durations, which may encompass months, years, or an entire lifetime ^63^.

In the current study, cerebral organoids in the pillar/perfusion plate were exposed to the model compounds for 1 and 3 weeks to investigate the effect of chronic exposure of compound on brain development. Ascorbic acid was used as a DNT negative control, which is frequently used as a supplement in various brain organoid differentiation protocols ^16,37,64–67^. It promotes the expansion of neuroepithelial-like stem cells by stimulating collagen synthesis through the MEK-ERK1/2 pathway ^68^. Methylmercury was selected as a DNT positive compound, which has been extensively studied for its acute toxicity ^11,12,69,70^. In our study, cerebral organoids were exposed to methylmercury starting from day 9, as this represents the initiation of neuroectoderm formation. Along the period of compound exposure, we monitored the changes in organoid morphology and the expression levels of neural marker genes. Our findings revealed that prolonged exposure to methylmercury resulted in the overexpression of the autistic *FOXG1* gene and a reduction in the organoid size. The underlying mechanism behind the *FOXG1* overexpression remains unknown, but it suggests that early exposure to methylmercury may induce autism spectrum disorder. While studying the exposure of DNT compounds to the brain, the existence of the proper blood-brain barrier is always a concern. Since our study has focused on investigating the influence of toxic compounds in the initial phase of brain development, there would be no need to incorporate the blood-brain barrier in cerebral organoid culture, considering its incomplete functionality in fetuses and newborns ^2–7^.

There are still areas for improvement in accurately assessing the impact of DNT compounds on the developing brain. Since DNT can occur at the late stage and post-birth as well, cerebral organoid culture needs to be maintained for longer time to attain full maturity and to assess late-stage DNT potential. In our study, cerebral organoids were cultured in the pillar/perfusion plate for up to 2 months, which might be insufficient to assess late-stage DNT. In addition, incorporation of vascularization and blood-brain barrier could provide more relevant data to predict the effects of DNT compounds at the later stage of brain development. While the pillar/perfusion plate could enable high-throughput assessment of DNT potential of compounds, the *in vitro* data obtained from cerebral organoids could not be correlated to the *in vivo* data directly. Thus, there is a need for sophisticated prediction models such as machine learning and AI with RNA sequencing datasets obtained from cerebral organoids and patients.

## Conclusions

Embryonic bodies (EBs), initially formed in an ultralow attachment (ULA) 384-well plate, were successfully transferred to the pillar plate containing Matrigel and subsequently differentiated into cerebral organoids in the perfusion plate. The dynamic culture of cerebral organoids in the pillar/perfusion plate significantly improved their growth and maturity compared to their static counterpart. The pillar/perfusion plate required only a small volume of cell culture media for 1 – 2 months of organoid differentiation, effectively alleviated the issue of diffusion limitation of nutrients and oxygen into the core of organoids *via* rapid, bidirectional flow on a digital rocker, and allowed *in situ* compound testing and organoid imaging. Day 9 cerebral organoids in the pillar/perfusion plate were exposed to ascorbic acid (DNT negative) and methylmercury (DNT positive) for 1 – 3 weeks to assess their DNT potential. We monitored changes in organoid morphology, size, and the expression levels of neural marker genes to evaluate the susceptibility of cerebral organoids to the compounds during differentiation. Cerebral organoids exposed to ascorbic acid exhibited no significant alterations in organoid morphology and gene expression whereas the organoids exposed to methylmercury displayed noticeable shrinkage in organoid size and abnormalities in neural gene expression. In summary, our results demonstrate that cerebral organoids cultured in the pillar/perfusion plate could be used for assessing the adverse effects of chronic exposure to toxic compounds on brain development.

## Acknowledgement

This study was supported by the National Institutes of Health (NCATS R44TR003491, NIDDK UH3DK119982, and NIEHS R43ES035653).

## Supplementary Information

**Supplementary Table 1.**
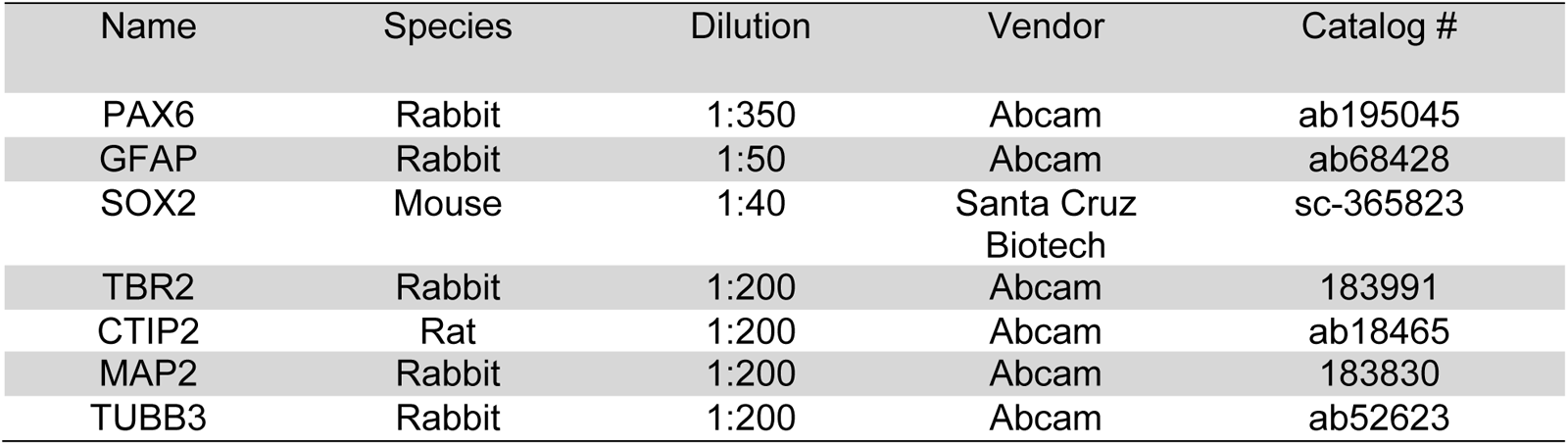
List of primary antibodies.

| Name | Species | Dilution | Vendor | Catalog # |
| --- | --- | --- | --- | --- |
| PAX6 | Rabbit | 1:350 | Abcam | ab195045 |
| GFAP | Rabbit | 1:50 | Abcam | ab68428 |
| SOX2 | Mouse | 1:40 | Santa Cruz Biotech | sc-365823 |
| TBR2 | Rabbit | 1:200 | Abcam | 183991 |
| CTIP2 | Rat | 1:200 | Abcam | ab18465 |
| MAP2 | Rabbit | 1:200 | Abcam | 183830 |
| TUBB3 | Rabbit | 1:200 | Abcam | ab52623 |

**Supplementary Table 2.** List of secondary antibodies.

| Name | Dilution | Vendor | Catalog # |
| --- | --- | --- | --- |
| Donkey anti-Rabbit IgG (H+L) Highly Cross-Adsorbed Secondary Antibody, Alexa Fluor 647 | 1:200 | ThermoFisher | A-31573 |
| Donkey anti-Mouse IgG (H+L) Highly Cross-Adsorbed Secondary Antibody, Alexa Fluor 488 | 1:200 | ThermoFisher | A-21202 |
| Donkey anti-Rabbit IgG (H+L) Highly Cross-Adsorbed Secondary Antibody, Alexa Fluor 488 | 1:200 | ThermoFisher | A21206 |
| Donkey anti-Rat IgG (H+L) Highly Cross-Adsorbed Secondary Antibody, Alexa Fluor 594 | 1:200 | Thermofisher | A-21209 |

**Supplementary Table 3.**
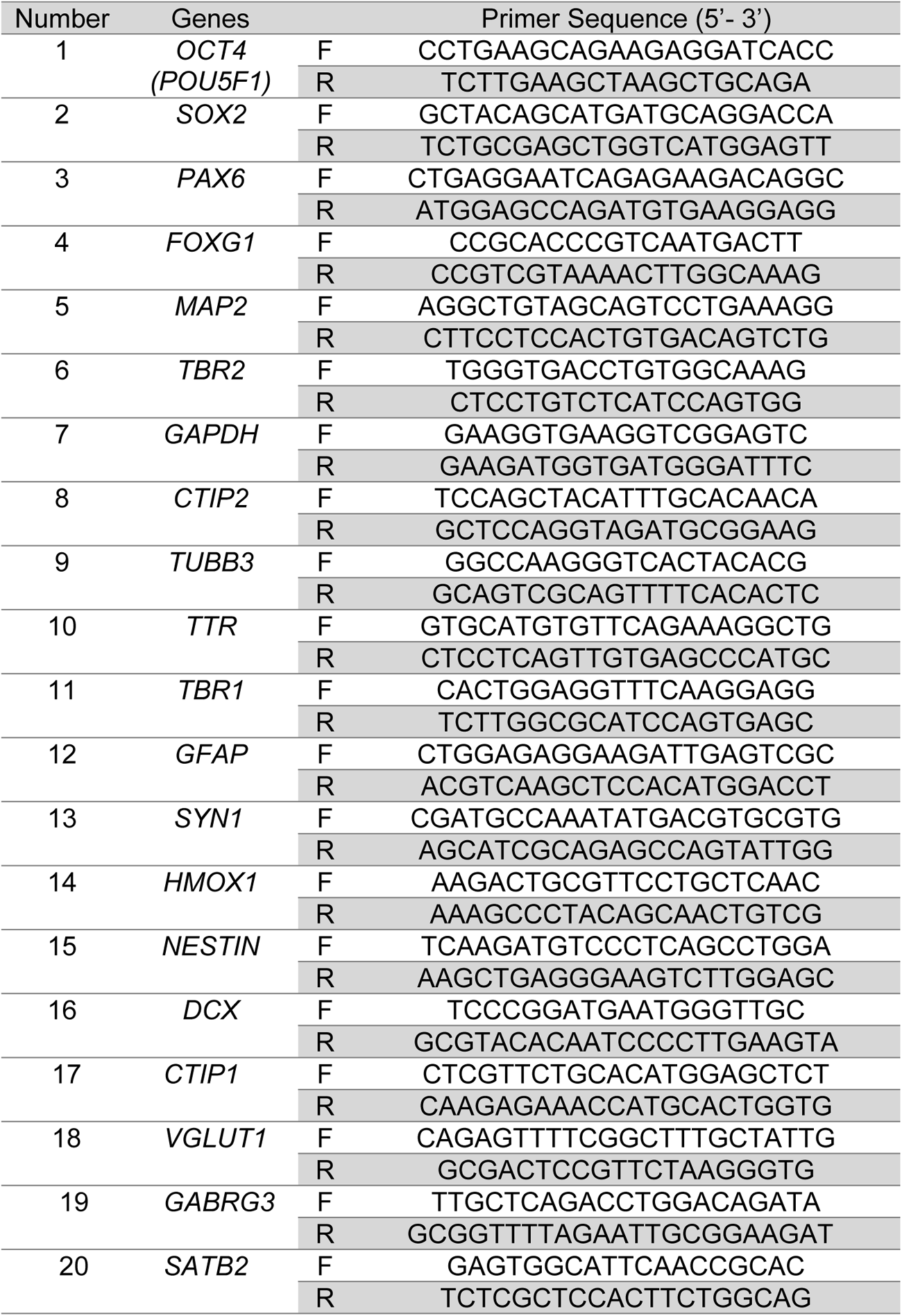
List of primers.

**Supplementary Figure 1.**
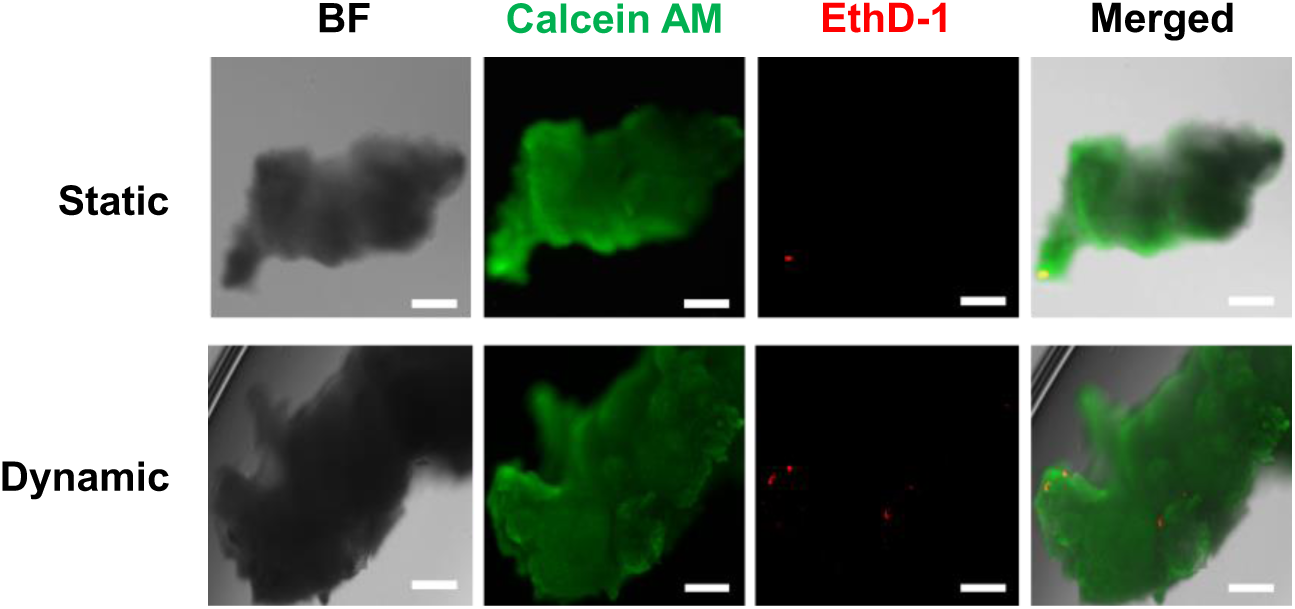
Measurement of cerebral organoid viability with calcein AM and ethidium homodimer (EthD-1) staining after static and dynamic culture. Scale bars: 200 µm.

**Supplementary Figure 2.**
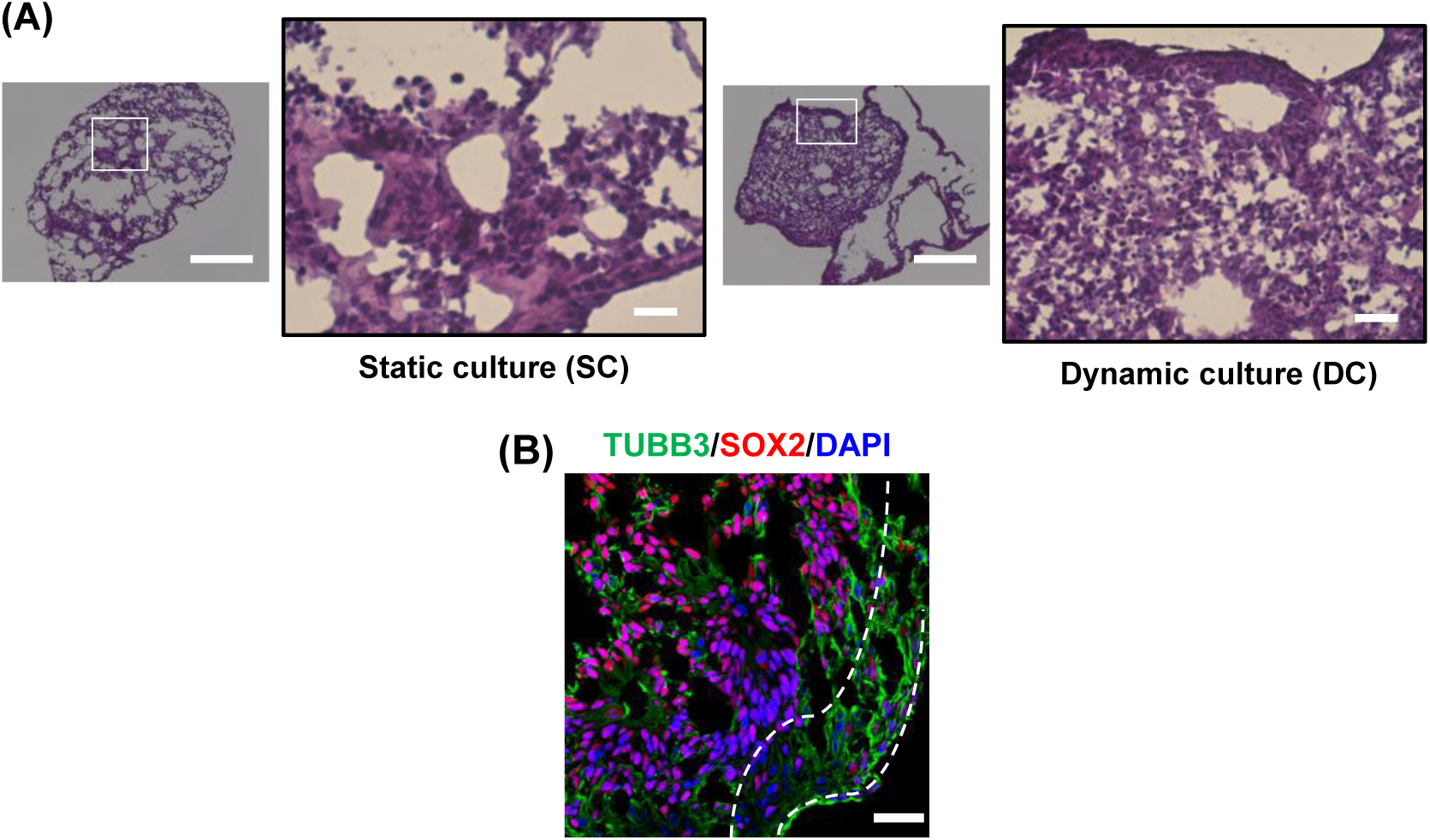
(A) H & E staining of day 31 cerebral organoids cultured in static and dynamic conditions. Dense cellular organization was observed in dynamically cultured cerebral organoids as compared to their static counterpart. The cavity structure in the H & E staining images is due to a very thin sectioning of the organoids (8 µm thickness) and the freezing artifacts. For cerebral organoids cultured in both static and dynamic conditions in the pillar/perfusion plate, we have not observed any cavity structures from bright-field imaging as well as whole-organoid immunofluorescence staining. **(B)** Immunofluorescence staining of cryo-sectioned cerebral organoids cultured in a dynamic condition for 31 days. Scale bar: 50 µm. The layering process was initiated in the dynamic-cultured cerebral organoids for day 31 with a spatial separation of *TUBB3* and *SOX2*.

**Supplementary Figure 3.**
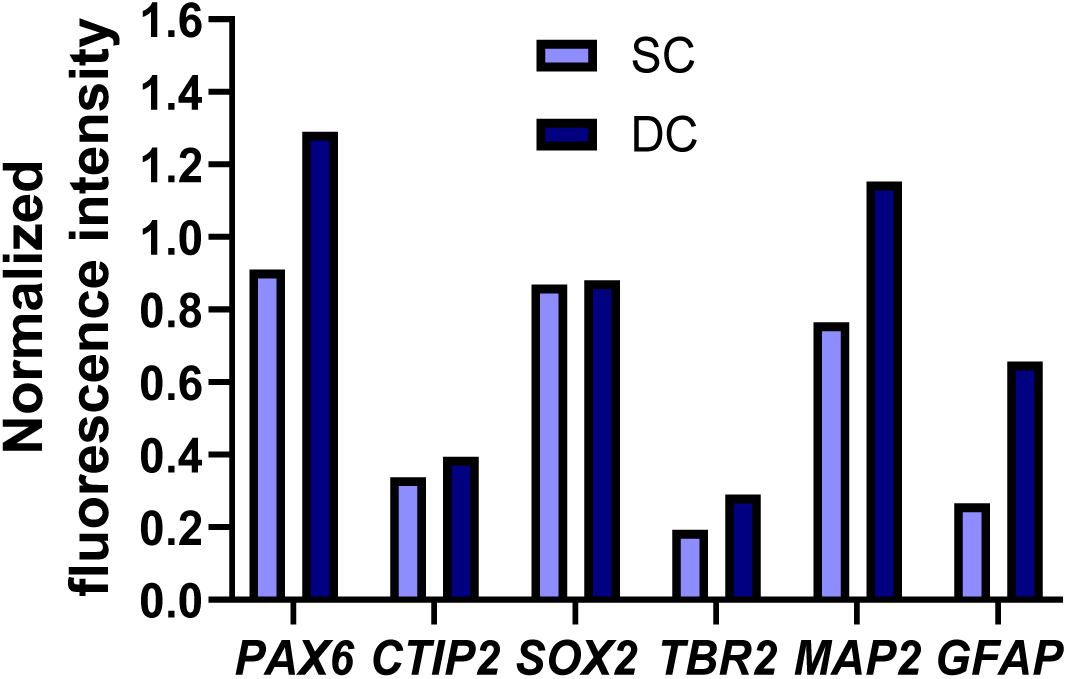
Normalized fluorescence intensity of cerebral organoids cultured in static and dynamic conditions. The fluorescence intensity of each biomarker was normalized with DAPI fluorescence intensity. SC represents static culture whereas DC represents dynamic culture of cerebral organoids.

**Supplementary Figure 4.**
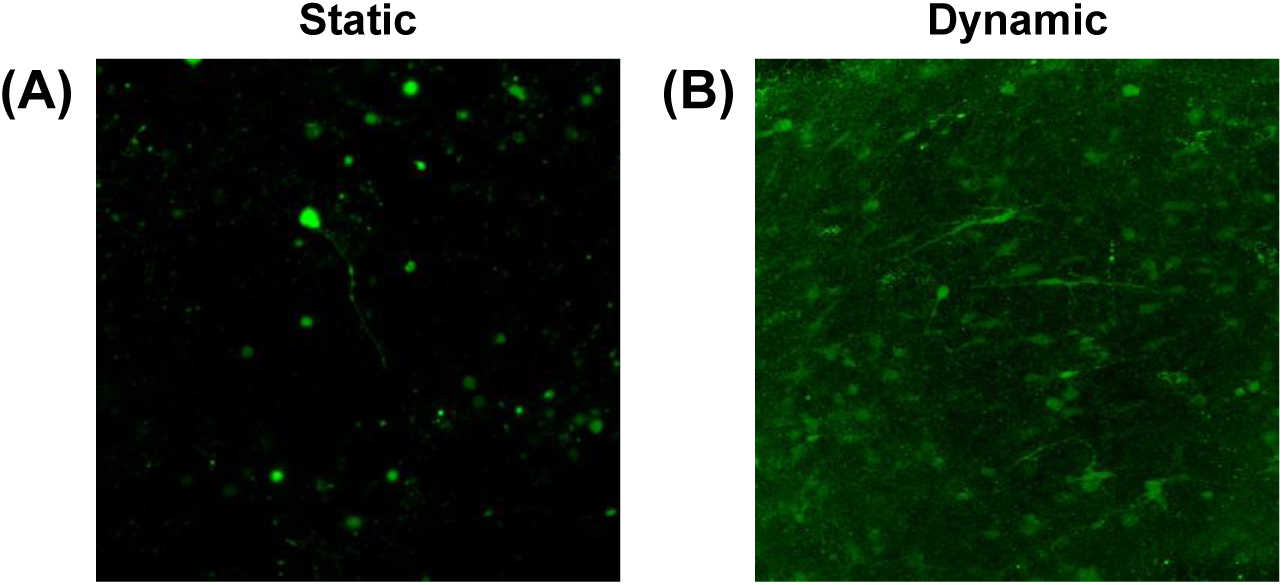
Calcium imaging of day 63 cerebral organoids cultured in **(A)** static and **(B)** dynamic conditions. Increased intracellular calcium ion was observed in dynamically cultured cerebral organoids as compared to the static counterpart, demonstrated by the increased green fluorescence.

**Supplementary Figure 5.**
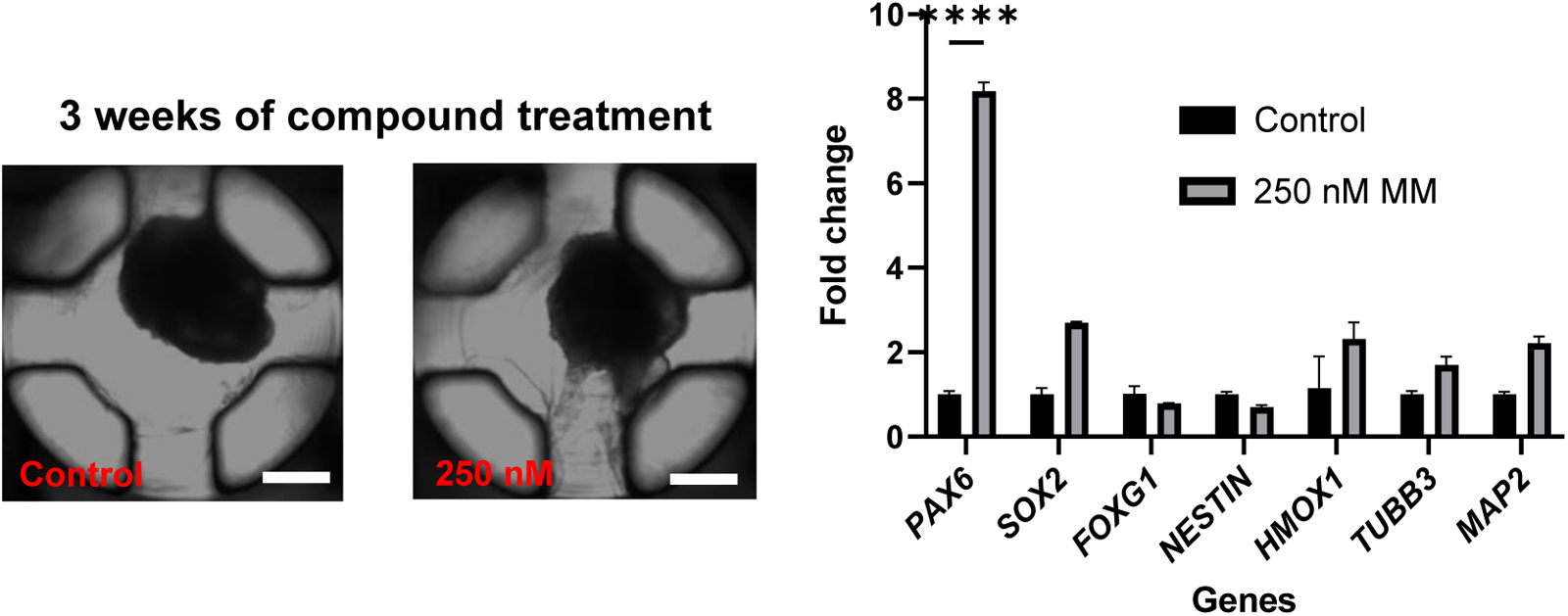
Cerebral organoids exposed to 250 nM methylmercury for 3 weeks in a static culture condition. Magnification: 4X, scale bars: 600 µm. No significant change in morphology was observed after 3 weeks of methylmercury treatment. The RT-qPCR analysis showed that *PAX6* neural progenitor marker was overexpressed when the organoids were exposed to 250 nM methylmercury. n = 12.

